# Genetic architecture of cichlid brain morphology

**DOI:** 10.64898/2026.04.01.715931

**Authors:** Jake Morris, David F. Rivas-Sánchez, Joel Elkin, Aaron Hickey, Bettina Fischer, Aleksandra Marconi, Richard Durbin, George F. Turner, M. Emilia Santos, Stephen H Montgomery

## Abstract

How evolutionary and developmental processes interact to determine axes of neural variation that produce behavioural diversity has been debated for many decades, with alternative hypotheses giving differential emphasis to functional coupling, which favours co-evolution, and developmental constraint, which enforces it. A critical omission is data on the genetic architecture of brain size and structure, which more closely illuminates the shared developmental dependencies between components of an integrated system. Here, we exploit ecological divergence between *Astatotilapia calliptera* and *Aulonocara stuartgranti*, two closely related cichlid species from Lake Malawi, to explore the genetic architecture of brain evolution. Using computer vision and machine learning techniques to extract volumetric data from micro-tomographic images, we first demonstrate significant divergence in brain composition between these species. Genomic and micro-tomographic imaging data from a population of hybrids generated between the two species were used to investigate genetic factors shaping this differentiation. We show that the majority of brain components are integrated phenotypically in hybrids, but genetic correlations between them are generally weaker. We further show that variation in multiple brain components is associated with variation in largely structure-specific quantitative trait loci, rather than determined by genetic factors with broad effects across the entire brain. These results suggest a genetic architecture that can facilitate modular changes in brain structure, and imply that individual components are independently evolvable.

## Introduction

The response of complex, composite traits to selection is, in part, determined by intrinsic factors that shape the permissible range of phenotypic variation (Arnold, 1992). Intrinsic constraints, including pleiotropy, can alter the path of evolution by influencing the availability of standing variation, and the fitness consequences of genetic variation across different aspects of trait development (Zhang, 2023). One context in which the relative roles of alternative sources of constraint have clouded our understanding of phenotypic variation, surrounds debates about how brains evolve. Brains are complex networks of neural circuits, which collectively optimise the behavioural response to external stimuli by processing environmental information in concert with an individual’s internal physiological state (Sterling & Laughlin, 2015). Brain size and structure are therefore expected to be intimately related to an animal’s ecology, but we have little resolution of how constraining forces that channel the evolution of brain structure and function could affect the direction or speed of ecological or behavioural divergence.

The conflicting roles of selection and constraint in shaping vertebrate brains have been debated for decades (Montgomery et al., 2016). Central to this debate are two hypotheses that have differing views on the origins and balance of constraining factors affecting brain evolution. Under one hypothesis, brain components are developmentally linked to overall brain size, such that the size of each component is largely determined by common developmental mechanisms, such as the schedule and timing of neurogenesis (Finlay & Darlington, 1995; Finlay et al., 2001). This would lead to major brain structures evolving in a coordinated, or concerted, way, with the size of separate components being closely predicted by total brain size. A contrasting hypothesis instead argues that variation in brain components is largely developmentally independent of both other brain structures, and of total brain size, allowing them to respond to targeted selection pressures in a more independent way, such that overall brain structure reflects a mosaic of responses to selection (Barton & Harvey, 2000). Mosaic evolution is often implicated in neural adaptation, reflected in non-allometric changes in brain structure, but mosaic models of brain evolution also invoke stabilising selection to maintain scaling relationships between co-evolving, functionally interdependent brain components (Barton & Harvey, 2000; Montgomery et al., 2016; Avin et al., 2021). A number of empirical findings closely align with the predictions of mosaic evolution, with units of functionally linked structures coevolving independently of more functionally disparate structures (Barton & Harvey, 2000; Gonzalez-Voyer et al., 2009a; Iwaniuk et al., 2004). However, other results hint at the influence of regulatory mechanisms with genetic effects shared across the entire brain, consistent with developmentally determined concerted evolution (Bond et al., 2002; Wang et al., 2008). Developmental data can therefore be invoked to support both hypotheses, but it is scarce relative to data on adult brain structure.

Distinctions between these ideas have become more nuanced, but universally satisfactory tests of their predictions remain elusive. This is, in part, due to the reliance of correlations between phenotypic variation in brain structures as a test of the prevalence of allometric, and non-allometric effects (Montgomery et al., 2016). This is problematic because patterns of coordinated evolution can emerge both through genetic constraint (as per “concerted” models) and strong functional interdependence (as per “mosaic” models) and are, as such, indeterminate when assessing the importance of developmental constraint on brain evolution (Avin et al., 2021).

To understand how brains evolve, greater emphasis is needed on the genetic causes of variation and their developmental effects, as concerted and mosaic models are, in effect, different visions of the genetic architecture of brain structure (Montgomery et al., 2016). Currently, there are limited studies investigating the quantitative genetics of brain evolution in natural systems. The examples that do exist largely report a high degree of genetic independence between brain structures (Hager et al., 2012; Henriksen et al., 2016; Noreikiene et al., 2015), with little evidence that patterns of phenotypic covariance are reflected by genetic covariance (Noreikiene et al., 2015). For example, using genetic mapping in a domesticated mouse strain, Hager et al. (2012) identified independent loci with isolated effects on individual brain regions. Using a similar approach in domestic chickens and their wild ancestors, Henriksen et al. (2016) showed independent genetic architectures for brain components and between brain and body size, while Noreikiene et al. (2015) found that genetic correlations between different brain components are generally low compared to corresponding phenotypic correlations in the three-spined stickleback (*Gasterosteus aculeatus*). These results therefore imply a distributed genetic architecture of brain structure, without strong genetic correlations between components that would otherwise limit the phenotypic response to selection on individual structures. However, because the data needed to answer these questions often requires the production of hybrid offspring to map quantitative traits, which can be costly and challenging to generate, these studies mostly concern standing genetic variation within domesticated populations (Hager et al., 2012; Noreikiene et al., 2015). Much of this genetic variation may be mildly deleterious, and as such it is possible selection for changes in brain structure may frequently act on *de novo* mutations distinct in their developmental effects from standing genetic variation (Avin et al., 2021; Montgomery et al., 2016). Therefore, data from additional natural systems representing more pronounced evolutionary divergence would be invaluable to conclusively investigate this issue.

Studies of natural radiations of closely related but ecologically diverse species, can provide insights into the genetic and developmental constraints that influence how neural systems respond to selection. During an adaptive radiation, multiple lineages proliferate within short timeframes into diverse ecological forms (Lamichhaney et al., 2015; Salzburger et al., 2014; Stroud & Losos, 2016). Local adaptation enables the exploitation of ecological opportunity (Schluter & McPhail, 1992) and promotes reproductive isolation among populations that increasingly differentiate phenotypically (Schluter, 2009), yet in many cases of adaptive radiation, species with divergent traits are able to produce viable hybrids (Rundle & Nosil, 2005). This implies that selection, rather than genetic incompatibility maintains population divergence, but also presents rich opportunities to understand the genetic architecture of divergent traits using interspecific crosses and the tools of quantitative genetics (Meredith, 1984; Pemberton, 2008).

Indeed, comparative studies have found striking diversity in sensory systems across closely related species, or ecotypes, in a variety of taxa with different ecological requirements and cognitive challenges (e.g. Catania, 2005; Huber et al.,1997; Hutcheon et al., 2002; Kotrschal & Palzenberger, 1992). For example, Mexican cave fish (*Astyanax mexicanus*) exhibit an enlarged hypothalamic region and smaller optic lobes compared to their ancestral surface-dwelling relatives (Loomis et al., 2019) and three-spined sticklebacks from benthic environments have distinct brain structures compared to limnetic ecotypes (Keagy et al., 2017). Both instances reflect heritable differences that correlate with the type of sensory information that is locally available. In radiations of African Great Lake cichlids, variation in brain morphology is pronounced, and linked to multiple ecological traits (Gonzalez-Voyer et al., 2009b; Huber et al., 1997; Pollen et al., 2007; Shumway, 2010; York et al., 2019). These differences in cichlid brain structure originate during early neural tissue patterning, attributable to variation in gene expression networks across species (Sylvester et al., 2010).

Here, we exploit emerging computer vision and machine learning techniques to provide new insights into the genetic architecture of brain evolution during an adaptive divergence. We developed an analytical pipeline to automatically segment and measure the volumes of major brain structures in two species of cichlid fish from Lake Malawi in Africa, and their interspecific hybrids, from CT-images. Our focal species, *Astatotilapia calliptera* and *Aulonocara stuartgranti*, belong to an extensive adaptive radiation (Loh et al., 2008; Malinsky et al., 2018), with *A. calliptera* a visual-hunting omnivore generally occupying shallow weedy areas, while *A. stuartgranti* uses its expanded lateral line system to detect prey hidden in patches of sediment in rocky habitats (Schwalbe et al., 2012). We first establish that brain size and structure of these species has diverged, consistent with the contrasting sensory environments of their foraging strategies. This then offered a unique opportunity to investigate the genetic architecture of brain structure by testing predictions of mosaic and concerted models while avoiding uncertainties related to the representativity of genetic variation within artificial populations.

## Materials and Methods

### Cross design and linkage map

We established a cross between a male from an *A. stuartgranti* population originally collected from near Usisya, Malawi, and two females from a *A. calliptera* population originally collected from the Itupi river in Southern Tanzania: the F1 offspring were interbred to produce F2 and F3 individuals. For the present study, only adult males were collected: a total of nine F_1,_ 129 F_2_ i150 F_3_. As males began to exhibit secondary sexual characteristics, they were isolated until they had grown to approximately 10cm standard length, and then l euthanised in 0.1% MS-222 solution, and fin clips were stored in 100% ethanol at-20°C prior to sequencing. DNA extraction and whole-genome sequencing library preparation was performed by the Wellcome Sanger Institute using an automated pipeline, and individuals were sequenced on NovaSeq 6000 S4 and NovaSeq X 10B platforms to an average depth of 4.5x – 8.6x for F_1_ to F_3_ samples. The raw sequences are available at NCBI BioProject PRJEB48145. Sample sequences were aligned to the fAulStu2 reference genome using BWA-MEM and converted to CRAM format using SAMtools *sort* (Danecek et al., 2021; Li, 2013). Genotype likelihoods were calculated using BCFtools *mpileup* and variants were called using BCFtools *call* (multiallelic model). For quality control, we filtered out sites with site depth outside of the range of the median +/- 25%, base quality below 20, median genotype quality below 35, or sample missingness above 50%.

To generate a linkage map for the cross, we started with 11,618 positions that are both alternately fixed between the parental populations, as well as heterozygous in all nine F_1_ offspring. Map construction was implemented with the R package OneMap (Margarido et al., 2007). F_2_ genotypes for all 11618 positions were binned (*find_bins* function) to eliminate uninformative positions for which all samples had the same genotype. Positions that exhibited significant distortion from Mendelian segregation were also filtered out (*test_segregation* function). Based on genotype probabilities, linkage groups were identified and markers were ordered along them according to recombination fraction using the unidirectional growth algorithm (*rf_2pts*, *make_seq*, and *ug* functions). Linkage groups corresponded correctly to the 22 chromosomes known in Lake Malawi cichlids.

### Diffusible iodine-based contrast-enhanced computed tomography

We used diffusible iodine-based contrast-enhanced computed tomography (Dice-CT) to visualise soft tissue and major brain structures. Each fish was stained for a minimum of 10 days in a 1% iodine, ethanol solution. This solution was refreshed every ∼7 days, after which the samples were washed in 100% ethanol for 30 minutes and then placed into 70% ethanol prior to imaging. The samples were imaged usomh a Nikon XTH 225ST X-ray tomgraphic scanner (Nikon Metrology, Tring, UK) with a resolution of approximately 30µm, an exposure of 708ms, 3141 projections and 1 frame per projection. 16-bit 3D image stacks were reconstructed using Nikon μCT software and VG Studio Max.

### Brain segmentation using machine learning

#### i. Manual segmentation of a training dataset generation and normalisation

Initial manual segmentations were performed in Dragonfly, applying a Contrast Limited Adaptive Histogram Equalization (CLAHE) technique to increase contrast (Zuiderveld, 1994). Six brain structures were segmented: the diencephalon, rhombencephalon, optic tectum, telencephalon, olfactory bulbs and brain stem. The posterior end of the brain stem was defined as 5 sections beyond the point at which the cross section of the brain stem closes into a consistent circular shape. In total, 75 fish, including 20 F2/F3s, ∼25 *Astatotilapia calliptera*, ∼15 *Aulonocara stuartgranti,* and 15 samples from the outgroup species *Rhamphochromis* sp. ‘chilingali’, were manually segmented. These segmentations were subsequently used as a training dataset for a deep-learning neural network model.

Prior to use as training data, each image was put through a pre-processing and normalisation pipeline that enhances and standardises the data by removing some variation in contrast, brightness and artefacts, including salt and pepper sign, between scans. This was carried out so that all samples were matched to a specific target scan of a sample that showed good contrast and was of approximately average size. The first step in the pipeline carried out a CLAHE using *cv2.apply.clahe()* from the python package *cv2* (*clip_limit=2.0*), before conversion to 8bit images using *cv2.normalise()*, and then applying a bilateral filter using *cv2.bilateralFilter()* to blur images while maintaining edges. We then made binary (approximate) masks of these images to separate the image into background vs foreground (the fish). This was done using the *cv2* functions *cv2.GaussianBlur()* with a kernel size of (7, 7)*, cv2.threshold()* using (cv2.THRESH_BINARY + cv2.THRESH_OTSU) and finally *cv2.morphologyEx()* first opening with cv2.MORPH_OPEN with a kernel of (3, 3) and then closing with cv2.MORPH_CLOSE with a kernel of (12, 12). This mask was then saved and used for histogram matching using the *match_histograms()* function from the *skimage* package (REF). Only the foreground (the fish) was histogram matched to avoid biasing this process in samples where more or less background was included due to size differences between samples. The final step in our pre-processing pipeline was a Z score normalisation. This was carried out by loading all 3D CT stacks and binary masks, and then calculating the global mean and standard deviation from all foreground pixel values. We then normalised all foreground pixels ((pixel - mean) / std) and then rescaled to the global max or 0-255 (for 8 bit images). Again, background pixels were left un-normalised. These Z score statistics were then saved so that they could later be applied to our target data.

#### ii. Core Model architecture

We used the python package Tensorflow/Keras (Abadi et al., 2016) to train 2.5D Unet models for automated, semantic segmentation of our CT scans. Briefly, Unets employ an encoder-decoder architecture with skip connections to learn to accurately identify features within images and their location. In our 2.5D Unet, the four encoder blocks were composed of [8, 16, 32, 64] filters while spatial dimensions were reduced by half at each layer using MaxPool2D(). These were then followed by a bottleneck of 128 filters, and four decoder blocks of [64, 32, 16, 8] filters with spatial dimensions doubling using UpSampling2D().

Each block consisted of two convolutional Conv2D() layers with k = (3, 3), and with ReLU activation to introduce non-linearity into the model to help it learn complex patterns.

In all blocks dropout was used for regularization, this helps prevent overfitting of the model to the training data. Finally, an output block with 1×1 convolution and sigmoid/softmax activation was used for pixel-wise classification, while we also implemented a resizing layer at the start of the process, making it possible to resize our input images. This provides two advantages, first it allows images with an odd number of pixels to be used (e.g out 401 x 401 cropped images) and two allows images to be made smaller, reducing computational power and increasing training speed.

#### iii. Image loading and augmentation

To maximise and balance computation efficiency while also harnessing some of the 3D context that is available in CT scans, we loaded images in such a way as to make our Unet 2.5D. This is done by collapsing slices into multi-channel images. In our case we used a total of 5 slices, one central slice (the target for segmentation), plus two above and two below (used for 3D context). Images were processed as batches, with this batch size customisable so that computation resources could be more easily managed. In addition, batches were loaded dynamically from disk rather than the alternative of loading the entire dataset into memory, this memory efficient design allowed model training in a more RAM limited environment. We also rescaled our input data using rescale=1./255, and implemented data augmentation to expand the size of our training dataset. This augmentation was carried out in sync so that geometric transformations (Rotation (180°), shearing (0.2), translation (0.2), and horizontal flipping) were applied to both images and manually segmented masks using matching random seeds. The number of rounds of augmentation was able to be set, but we typically applied five rounds to our dataset, increasing the size of training set from ∼80 individual scans, to a total of ∼480 including the augmented and original scans.

#### iv. Model training

In each round of training, models were trained for 150 epochs. The first 2500 steps were used for warmup with the learning rate starting at 0 and increasing untill it hit the desired starting learning rate (after approximately the first two epochs), before it declined over the next 100 epochs using cosine decay. This slow decline in learning rate allows the model to first make large jumps in learning, before making smaller more refined adjustments as the model improves in later epochs. We used weighted categorical cross entropy as our loss function during training, and the generalised dice coefficient as our validation metric.

These functions have the advantage of weighting target classes based on their representation in the training data, so that smaller structures have a greater effect on model optimisation than larger classes. This is important where class imbalances are large, as it prevents for example the model being given a score of 0.95 by simply predicting everything incorrectly as background because 95% of the pixels in the dataset are background pixels. Approximately 20% of our training dataset was used for validation while the rest was used for training (although different subsets were used for training at times). We trained two models for segmentation of different structures, the first segmenting on the brain stem, rhombencephalon and diencephalon, and the second segmenting the diencephalon, optic tectum and telencephalon.

#### v. Inference and segmentation

We first applied our normalisation pipeline to our target dataset (the rest of our dataset), using histogram matching to the same sample as for the training dataset, and using the global mean and standard deviation from our training set, so that our target samples matched our training data more closely. We then used Tensorflow to apply our final trained model to our target data and make predictions for each pixel. Briefly, this was done by loading in each 2.5D image stack (so that 3D context is used during inference) and then producing probability maps from our trained model for each pixel class, before using argmax to convert probabilities to class predictions. This results in a mask showing the most probable class for each pixel in each slice of our sample stack.

We then used a custom script to automatically load our data into Dragonfly (ORS), and apply some standard functions to this data to produce regions of interest (ROIs) for manual correction. In brief, the script first loads all three image stacks, the normalised scans, and the two sets of masks produced by each of our two models (one brain stem, rhombencephalon and diencephalon, and the other Diencephalon, optic tectum, and Telencephalon). It then uses the ORS function getAsROIWithinRangeInArea() to convert matching pixels from our masks into 3D ROI coordinates, so that class 1 from our first model becomes our brain stem ROI, class 2: from our first model becomes our rhombencephalon, class 3 from our first model becomes our diencephalon ROI, class 1: from our second model becomes our optic tectum ROI, and class 2: from our second model becomes our telencephalon ROI. We then applied several morphological operations to these ROI, first removing all islands <750 pixels; then smoothing using kernelShape =’square’, kernelDim = 3, kernelSize = 9; followed by erode using kernelShape =’square’, kernelDim = 3, kernelSize = 3; and finally dilate using kernelShape =’square’, kernelDim = 3, kernelSize = 3. After this we manually inspected and refined these ROI using the original stack to confirm they were accurately segmented. Due to the small size of the olfactory bulb we segmented this structure manually in all samples.

### Differences among parentals and hybrids

We used the *A. calliptera* (n=25) and *A. stuartgranti* (n=15) individuals that we manually segmented to build our training data set to investigate brain structure differences between the parental species. For these analyses, we applied a logarithmic transformation to volumetric measurements of the brain structures and used the brain stem as an allometric control, following confirmation that its volume does not differ significantly between the species (*F*_1,38_ = 2.389, p = 0.130). We then used linear models to assess variation in the volume for each structure of interest (namely the rhombencephalon, the optic tectum, the diencephalon, the telencephalon or the olfactory bulb) as a function of the brain stem and species. We determined the importance of the independent variable “species” by comparing the residual sum of squares (RSS) of models, with and without the term, as well as implementing Wald 𝜒^2^ tests with the *Anova* function of the R package *CAR* (Fox & Weisberg, 2018).

We then explored how brain structures in hybrids (n=275) vary relative to the two parental species and investigated the nature of volumetric differences among hybrids and their parentals. This was done by first, conducting a principal component analysis (PCA) and testing for group-differences on major principal components (PCs) via Kruskal-Wallis and post-hoc Dunn’s tests (Benjamini-Hochberg p-adjustment method). This was followed by standardised major axis regression analyses (*SMATR* package in R (Warton et al., 2012)) incorporating the allometric equation log *y*=*β*log *x* + *α*, which describes scaling between a trait (*y*) and an allometric control (x). In these analyses, differences between species are stablished by comparing the slope (*β*) and the intercept (*α*) of their corresponding allometric lines. Changes in slope suggest distinct scaling relationships across groups whereas differences in the intercept indicate non-allometric grade-shifts (Kruska, 2005), both of these instances indicate non-allometric change in a brain trait, attributable to selection (Barton & Harvey, 2000).

### Estimating genetic correlations

To identify the strength of genetic covariance between pairwise brain structures, we calculated 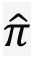 between pairs using the PLINK genetic toolset (Chang et al., 2015). To this end, we first produced a VCF file (variant call format) with information on our samples (SNPs with DP > 25, Qual >20, GQ >30 and missingness ≤ 50%) using a snakemake pipeline incorporating bcftools. We then converted this VCF into a.bed file (genotype matrix) spanning chromosomes 1-23, alongside.bim and.fam files with complementary information. We then applied LD (linkage disequilibrium) pruning using a window size of 50 SNPs, a step size of 5 SNPs, and a r^2^ threshold of 0.2, to create a list of independent SNPs contained in a subsequent LD-pruned-bed file. This allowed us to compute 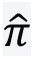 using PLINK’s *–genome* function, which makes pairwise comparisons between individuals to obtain IBD (identity by descent) measurements and build a relatedness matrix using a python script.

To calculate pairwise genetic correlations between brain structures, we used a python script that implemented a bivariate animal model with PyMC (probabilistic programming library), including brain stem as a covariate. Our model used four chains and 8,000 tuning steps followed by 5,000 sampling draws. The model decomposed each trait as genetic and residual effects with brain stem as a fixed effect. Traits were standarised to mean=0 and SD=1. We used Cholesky decomposition of the relatedness matrix to obtain its square root, improving the speed and stability of the model. Genetic and residual 2×2 covariance matrices were parameterised using LKJCholeskyCov() priors, with η = 4.0 and a standard deviation scale of 0.7, placing moderate prior weight around zero correlation while setting expectations for genetic standard deviations. We included brain stem volume as a fixed covariate in the model and genetic, random effects were marginalised, avoiding explicitly sampling individual genetic values while maintaining the correct covariance structure. For each trait pair, this model outputs posterior estimates of genetic and residual phenotypic correlations, alongside 95% credible intervals derived from posterior genetic and residual phenotypic covariance matrices. A Mantel test (999 permutations) implemented with R (*mantel* function R package Vegan (Oksanen et al., 2001)) was used to test for a potential correspondence between our phenotypic and genetic correlation matrices.

### Quantitative Trait mapping analysis

We carried out a quantitative trait loci (QTL) analysis across the 23 autosomes using the R/qtl2 package in R (Broman et al., 2019). First, we assembled a genetic map with pseudo markers at 1cM intervals (function *insert_pseudomarkers*) and calculated the genotype probabilities at each position with an error rate of 0.002 (function *calc_genoprob*). Then, we computed a kinship matrix to account for relatedness between individuals (function *calc_kinship*). Next, for each trait, we carried out a QTL scan (function *scan1*) with linear models that consider relationships among individuals as a random polygenic effect (our kinship matrix) and brain stem volume as a covariate. This was followed by genome scans with 1000 permutations (function *scan1perm*) to calculate the significant thresholds of logarithm of the Odds (LOD) ratios and identify peaks in LOD curves (function *find_peaks*; levels of significance *α* = 0.05 and 0.1) associated with QTL positions exceeding these thresholds. Then, we estimated Bayesian credible intervals (95%, function *bayes_int*) for these positions. Fnally, to investigate the relative contribution of significant (*α* = 0.05) QTLs to trait variation, for each trait, we estimated heritability due to the QTL and converted it into a percentage of variance explained by the QTL (percentage = 1-10 ^-2/n^ ^LOD^).

## Results

### *i)* Extensive divergence in brain composition between *A. calliptera* and *A. stuartgranti*, with variable, intermediate traits in hybrids

Accounting for allometric effects, our linear models revealed significant differences in the volume of all brain structures between species (rhombencephalon, *F*_2,37_ = 21.66, p = <0.001; optic tectum, *F*_2,37_ = 67.23, p = <0.001; diencephalon, *F*_2,37_ = 96.42, p = <0.001; telencephalon, *F*_2,37_ = 24.71, p = <0.001 and olfactory bulb, *F*_2,37_ = 121.80, p = <0.001; Figure 1). A PCA on these structures produced two PCs that collectively explained 87.47% of the variation in brain structure in parental species and hybrids (Figure 2A). PCs were significantly different in all pair-wise comparisons involving the three groups (PC1, 76.44% Var, Kruskal-Wallis *χ*^2^_2_ = 66.079, *P* < 0.001, eigenvalue=3.822 and PC2, 11.02% Var, Kruskal-Wallis *χ*^2^_2_ = 37.801, *P* < 0.001, eigenvalue=0.551) and were consistent with hybrids having intermediate trait values (Figure 2B-F). Allometric scaling analyses confirmed this variation consists of non-allometric shifts in the slope (diencephalon, LR_2_^2^ = 15.890, *P* < 0.001; telencephalon, LR_2_^2^ = 18.72, *P* < 0.001 and olfactory bulb LR_2_^2^ = 16.37, *P* < 0.001) and the intercept (rhombencephalon, Wald *χ*^2^_1_ = 44.63, *P* < 0.001 and optic tectum Wald *χ*^2^_1_ = 147, *P* < 0.001) of the scaling relationships between brain components against the brain stem. This results in generally larger brain structures in *A. calliptera* relative to *A. stuartgranti*, but with variable effect sizes, and intermediate volumes in hybrids for a given brain size, despite steeper scaling in some *A. stuartgranti* structures (Figure 1A-E, Table 1).

**Figure 1.**
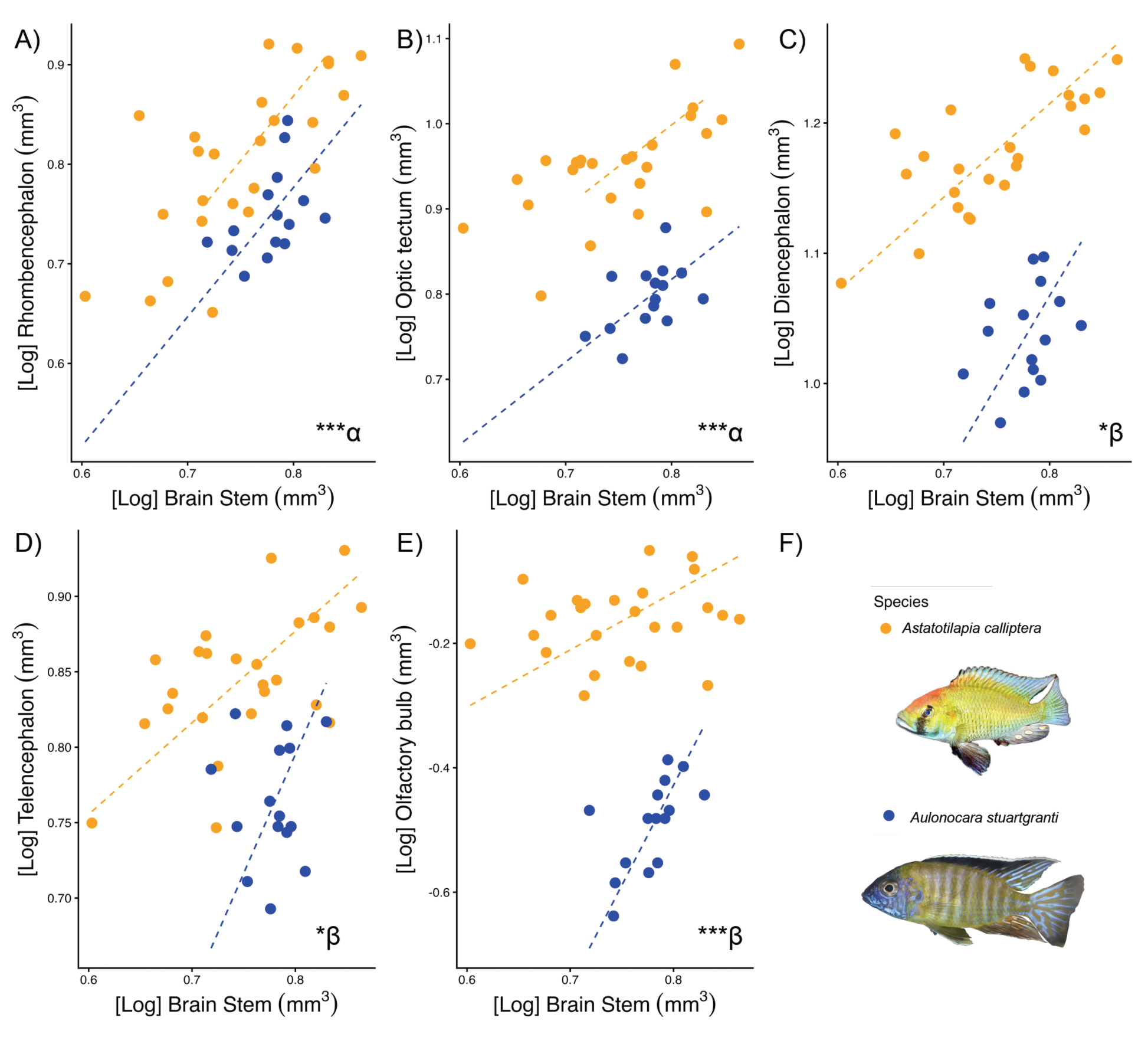
Inter-specific non-allometric shifts in scaling of the volume of the A) rhombencephalon, B) diencephalon, C) optic tectum, D) telencephalon and E) the olfactory bulb against an allometric control. Points and dashed lines represent respectively individuals and scaling relationships between brain components and the brain stem volume for *A. calliptera* (orange) and *A. stuartgranti* (blue). Asterisks convey significance levels of intercept (⍺, Wald 𝜒2 statistic) slope (*β*, LR) shifts. * p < 0.05. ** p < 0.01. *** p < 0.001.

**Figure 2.**
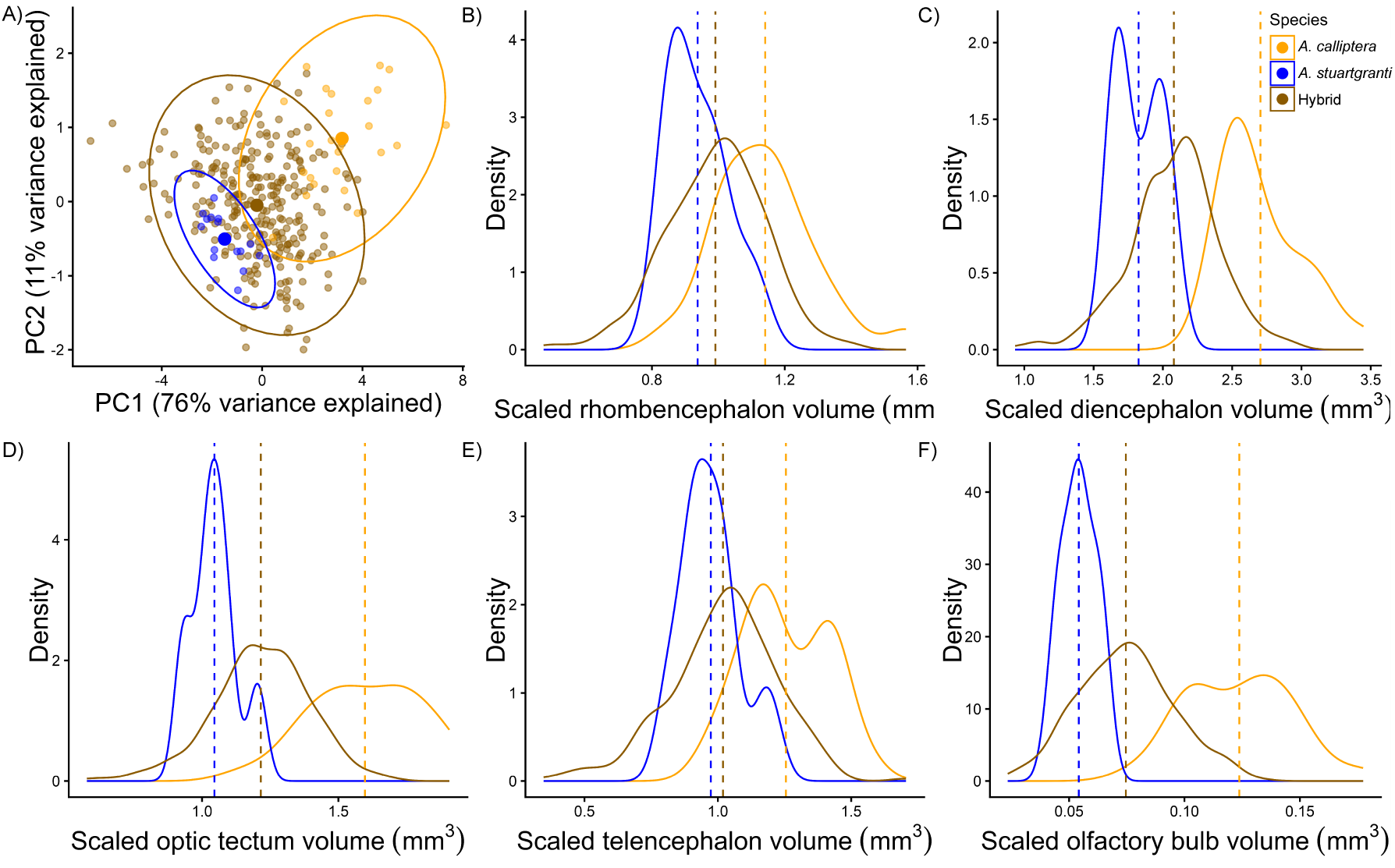
(A) Biplot between PC1 and PC2 from brain structure variation. (B-F) Density plots showing the distribution of volume variation of the B) rhombencephalon, C) diencephalon, D) optic tectum, E) telencephalon and F) the olfactory bulb scaled against the brain stem in *A. calliptera* (orange), *A. stuartgranti* (blue) and all hybrids between the two (brown).

**Table 1.** Results from SMATR regressions, showing likelihood ratio statistics, Wald scores, pair-wise statistics and P-values of slope and elevation shift tests.

| Slope ($\beta$) shift | | | | | |
| --- | --- | --- | --- | --- | --- |
| Brain component | LR<sub>2</sub> | P | Pair-wise comparison | T-statistic | P |
| Rhombencephalon | 0.689 | 0.708 | *A. calliptera*-*A. stuartgranti* | 0.402 | 0.526 |
|  |  |  | *A. calliptera*-Hybrid | 0.559 | 0.455 |
|  |  |  | *A. stuartgranti*-Hybrid | 0.111 | 0.739 |
| Optic tectum | 2.943 | 0.223 | *A. calliptera*-*A. stuartgranti* | 1.601 | 0.206 |
|  |  |  | *A. calliptera*-Hybrid | 2.684 | 0.101 |
|  |  |  | *A. stuartgranti*-Hybrid | 0.215 | 0.643 |
| Diencephalon | 15.890 | <0.001 | *A. calliptera*-*A. stuartgranti* | 4.918 | 0.027 |
|  |  |  | *A. calliptera*-Hybrid | 15.770 | <0.001 |
|  |  |  | *A. stuartgranti*-Hybrid | 0.043 | 0.837 |
| Telencephalon | 18.720 | <0.001 | *A. calliptera*-*A. stuartgranti* | 6.763 | 0.009 |
|  |  |  | *A. calliptera*-Hybrid | 18.671 | <0.001 |
|  |  |  | *A. stuartgranti*-Hybrid | 0.011 | 0.917 |
| Olfactory bulb | 16.370 | <0.001 | *A. calliptera*-*A. stuartgranti* | 16.31 | <0.001 |
|  |  |  | *A. calliptera*-Hybrid | 7.782 | 0.005 |
|  |  |  | *A. stuartgranti*-Hybrid | 7.710 | 0.005 |
| Elevation ($\alpha$) shift | | | | | |
| Brain component | Wald $\chi^2$ | P | Pair-wise comparison | T-statistic | P |
| Rhombencephalon | 44.630 | <0.001 | *A. calliptera*-*A. stuartgranti* | 27.153 | <0.001 |
|  |  |  | *A. calliptera*-Hybrid | 43.486 | <0.001 |
|  |  |  | *A. stuartgranti*-Hybrid | 0.365 | 0.546 |
| Optic tectum | 147.00 | <0.001 | *A. calliptera*-*A. stuartgranti* | 151.421 | <0.001 |
|  |  |  | *A. calliptera*-Hybrid | 89.331 | <0.001 |
|  |  |  | *A. stuartgranti*-Hybrid | 21.737 | <0.001 |

### ***ii)*** Genetic correlations between brain components suggest weaker ties than phenotypic covariance

With the exception of trait pairs that involve the olfactory bulb, the majority of the genetic correlations between brain components were weaker than the corresponding phenotypic covariances (Figure 3). The 95% Bayesian confidence intervals of some of our posterior means contain zero, however, the upper confidence intervals of these estimates are, for multiple trait pairs (diencephalon-rhombencephalon; telencephalon-rhombencephalon; optic tectum-diencephalon; telencephalon-diencephalon; optic tectum-telencephalon), near or below the corresponding posterior residual phenotypic covariance. A Mantel test comparing our genetic and phenotypic matrices showed a moderate, positive but not significant correspondance between them (Mantel Test: r = 0.488, p = 0.166), suggesting genetic correlations mirror, but do not closely predict phenotypic covariance.

**Figure 3.**
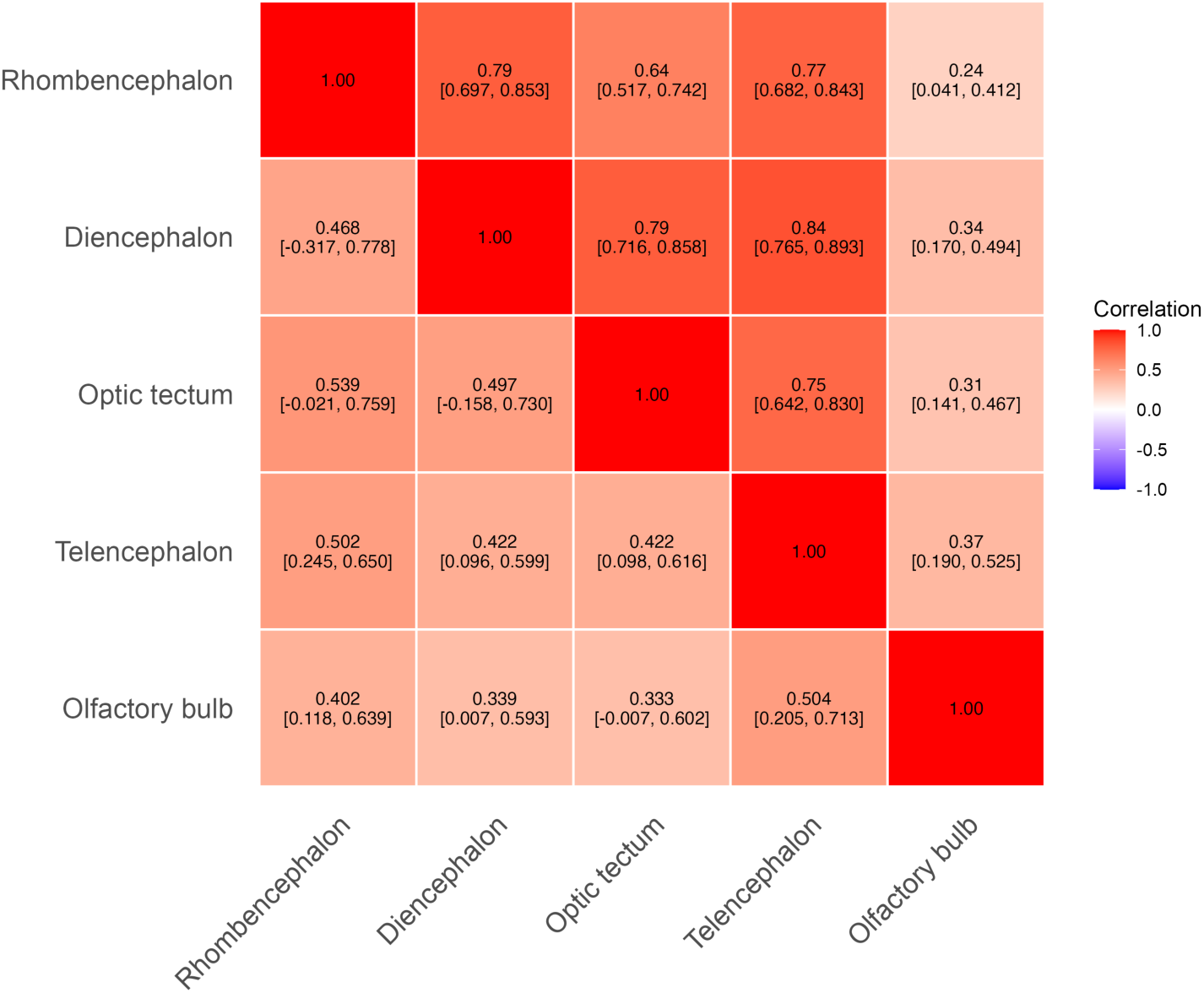
Heatmap of correlations of genetic and residual phenotypic correlations between higher brain structures, estimated from an animal bivariate model with brain stem as a fixed effect. Above the diagonal, posterior mean of residual (after genetic and brain stem effects are removed) phenotypic correlations. Below the diagonal, posterior mean of genetic correlation. For each tile, 95% Bayesian credible intervals in brackets.

### ***iii)*** Independent major QTL for individual brain structures

We identified three significant QTLs associated with volumetric variation in brain structure. One on chromosome 7, associated with the rhombencephalon (LOD = 4.061; Pos = 27cM; CI_l_ 5.888 - CI_u_ 50.365; 6.48% of trait variation explained; Figure 4A), a second on chromosome 13, associated with telencephalon volume (LOD = 4.881; Pos = 12.21cM; CI_l_ 3.170 - CI_u_ 14.189; 7.74% of trait variation explained; Figure 4B), and a third on chromosome 9, associated with the optic tectum (LOD = 4.370; Pos = 30.01cM; CI_l_ 29.316 - CI_u_ 68.673; 6.95% of trait variation explained; Figure 4 c). In addition, we identified putative (*α* = 0.1) QTLs on chromosomes 20 (LOD = 3.677; Pos = 29.86cM; CI_l_ 3.132 - CI_u_ 44.121; 5.88% of trait variation explained; Figure 4D) and 13 (LOD = 3.778; Pos = 11.22cM; CI_l_ 3.170 - CI_u_ 13.383; 6.04% of trait variation explained; Figure 4C), associated with variation in the olfactory bulb and optic tectum respectively, as well as a putative QTL on chromosome 13 associated with overall brain size (LOD = 3.633; Pos = 11.221; CI_l_ 3.170 - CI_u_ 30.465; 5.82% of trait variation explained; Figure 4F). We did not detect significant or tentative QTLs linked to diencephalon variation (Figure 4 e).

**Figure 4.**
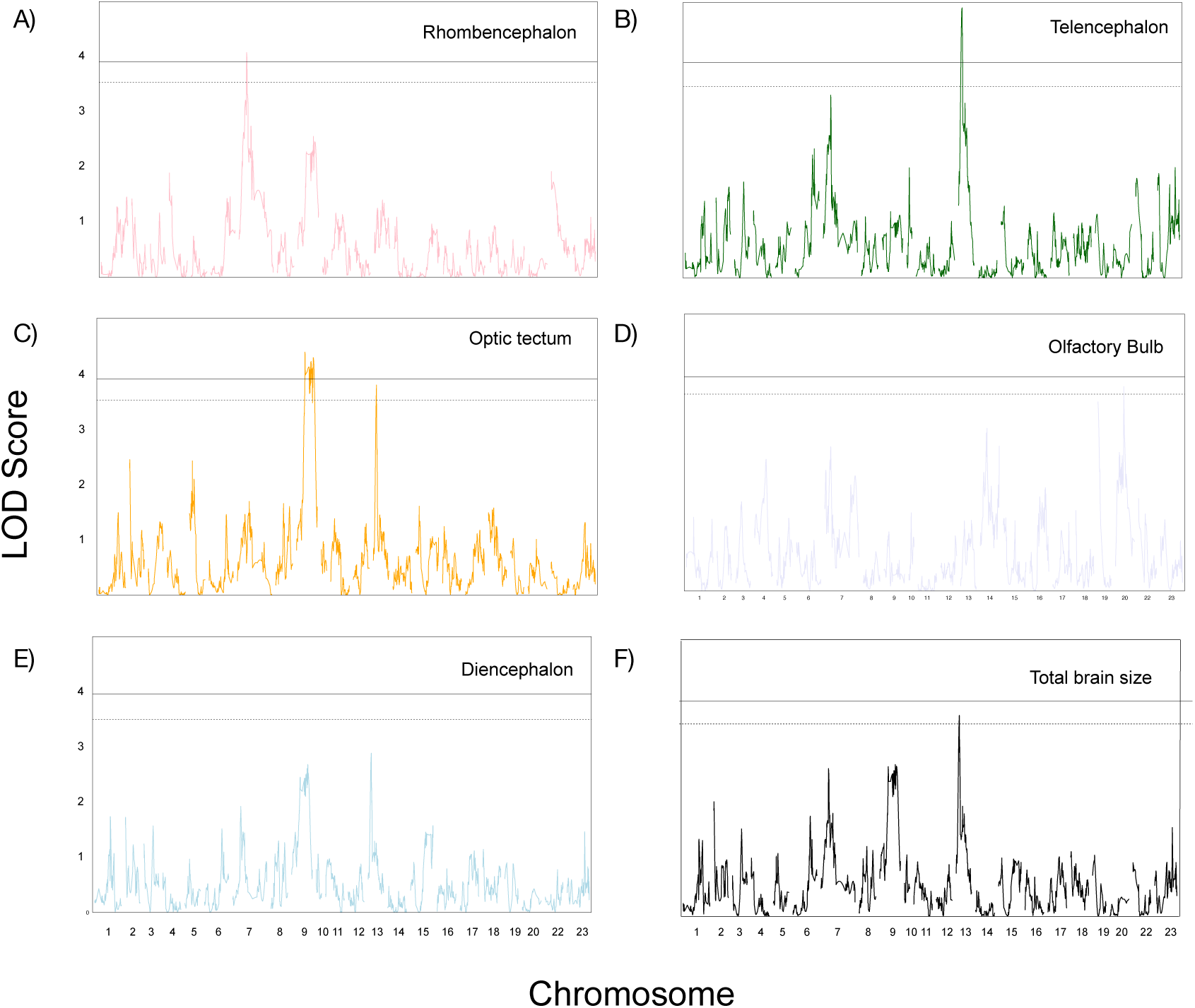
Quantitative trait loci (QTL) associated with variation in the volume of the rhombencephalon (a), the telencephalon (b), the optic tectum (c), the olfactory bulb (d), the diencephalon (e) and total brain size (f). In each plot, the dotted and solid horizontal lines represent, respectively, the LOD threshold of tentative (α<0.1) and significant (α<0.05) QTLs.

The positions of the significant telencephalon QTL, the tentative optic lobe QTL and the tentative brain size QTL on chromosome 13 are similar, potentially reflecting a shared genetic component among these traits. To test for a pleiotropy, in which a single QTL is associated with variation in multiple structures, we performed additional QTL analyses with the optic tectum as the focal trait, with and without the inclusion of the telencephalon as a covariate. When telencephalon variation was accounted for, the putative optic tectum QTL collapsed, consistent with a shared genetic effect across the two structures. We also investigated whether the brain size QTL on chromosome 13 reflects accumulated genetic effects on the telencephalon and the optic tectum by performing brain volume genome scans that exclude either of these structures as an explanatory term, with brain stem as a covariate. No QTL peaks above the LOD significance thresholds were detected in these scans, potentially indicating that the signal for total brain size is driven by the combined effects of a single QTL on chromosome 13 influencing the telencephalon and the optic tectum.

## Discussion

We investigated brain structure variation in two species of cichlid fishes from Lake Malawi that belong to a recent, prolific adaptive radiation. We demonstrated that *A. stuartgranti,* which occupies a low-light sensory environment, exhibits non-allometric reductions in multiple brain components relative to the generalist, *A. calliptera*, which occupies brighter habitats. While multiple brain components shift in a similar direction, the pattern and extent of divergence vary across brain regions, suggesting a complex, heterogenous pattern of divergence. Assisted by computer vision techniques and genomic data on a large sample of hybrids between the two species, we explored the genetic architecture underlying these patterns of divergence. We found that, with the exception of pairwise comparisons involving the olfactory bulb, genetic correlations between brain components are low relative to the corresponding phenotypic covariances. We also detected major quantitative trait loci associated with variation in the volume of the telencephalon, the optic tectum and the rhombencephalon. This results generally indicate a degree of genetic independence among brain structures, which supports an inference of a distributed genetic architecture, in line with mosaic models of brain evolution.

The mosaic brain hypothesis predicts that selective pressures on behaviour will be met by behavioural specialisations mediated by functionally linked brain components (Barton & Harvey, 2000). These adaptive responses involve selective changes in the level of neural investment on components underlying particular behavioural tasks. We found that *A. stuartgranti* has smaller brain components than *A. calliptera*, attributable to heritable, volumetric reductions that cannot be explained by allometric scaling with the rest of the brain, and may instead be associated with ecological variation. Species of the genus *Aulonocara* have widened cephalic lateral line canals that can identify hydrodynamic stimuli emitting from prey concealed in the sediment (Schwalbe et al., 2012). Many *Aulonocara* can therefore forage in dimly lit environments, active at dawn and dusk, or inside caves, with some species penetrating to depths of 90 m of more, at which visual prey detection is not possible for potential competitors, such as *A. calliptera* (Schwalbe & Webb, 2015). In contrast, *A. calliptera* is generally confined to well-lit waters where it visually locates food items (Joyce et al., 2011; Konings, 1989; Parsons et al., 2017).

Because *A. stuartgranti* uses visual cues to find prey facultatively rather than obligatorily (Schwalbe & Webb, 2015), selection for visual capacity in its dimmer sensory environment may be relaxed. Thus, the high costs associated with neural tissue investment (Laughlin, 2001; Laughlin et al., 1998; Niven & Laughlin, 2008) could outweigh the benefits of supporting an investment in visual processing, resulting in volumetric reductions of the optic tectum. Similar patterns of reduced visual investment associated with ecological divergence have been reported in Mexican cave fish populations where the optic tectum has regressed multiple times relative to the surface ecotype (Loomis et al., 2019), and in other Malawi cichlids that forage in turbid waters and which exhibit reduced optic tecta (Huber et al., 1997). However, we also found that *A. stuartgranti* has smaller olfactory bulbs compared to *A. calliptera* despite, in principle, olfaction and taste being sensory modalities that could be favoured in dark aquatic habitats such as those exploited by *A. stuartgranti* (Brandstätter & Kotrschal, 1990; Kotrschal & Palzenberger, 1992). Additional research is required to pinpoint specific mechanisms linking ecological divergence and additional sources of divergent selection that result in these non-allometric, volume reductions that also affect downstream brain components (diencephalon, the telencephalon and the rhombencephalon).

Modular changes in the brain are expected to be constrained by genetic correlations between components that limit the extent to which each can evolve independently (Finlay & Darlington, 1995). Exploring these correlations requires study designs often precluded in natural populations due to sample size restrictions (Moore & Kukuk, 2002). However, artificial neural network models have become available for categorising large image datasets (Mahmud et al., 2021), enabling large population sampling through relatively quick extraction of meaningful biological information (Lürig et al., 2021; Mahmud et al., 2021). We took advantage of this approach using deep-learning models to segment brain images of *A. calliptera* and *A. stuartgranti* hybrids, to provide a basis for quantitative genetics analyses. We found that, after accounting for allometric scaling, the majority of the phenotypic covariances between components are higher than the corresponding genetic correlations, which are in turn, generally closer to zero than to unity. The discrepancy between genetic and phenotypic covariance was the highest in pair-wise comparisons involving the diencephalon, the optic tectum, the telencephalon and the rhombencephalon. Although this, in part, may reflect uncertainty in the estimation of genetic correlation, the estimated strength of genetic correlations may imply that phenotypic data may produce false assessments of the strength of developmental integration (Noreikiene et al., 2015), and suggest that there are avenues through which these components can potentially respond to selection individually, despite a degree of phenotypic integration (Barton & Harvey, 2000).

These results are supported by our QTL analysis, in which we detected significant, independent, loci associated with the volume of the telencephalon, the optic tectum, and the rhombencephalon as well as a tentative QTL influencing the olfactory bulb. While these traits are all likely polygenic with many small effect loci, the observation that the largest effect loci detected in these QTLs are independent of overall brain size reinforces our interpretation of incomplete genetic correlations between structures. This suggests a genetic architecture by which neural divergence between *A. calliptera* and *A. stuartgranti* has been shaped by independent evolutionary responses to selection acting separately on different components. In contrast, no major loci for overall brain size were identified, and the strongest candidate QTL likely reflects the compound influence of two structures. This may in turn suggest selection on loci controlling brain structure shape the evolution of brain size, rather than the reverse. These findings mirror quantitative genetic analyses in other vertebrate systems, where brain components are rather weakly genetically correlated (Hager et al., 2012; Noreikiene et al., 2015), and where mosaic evolution is thought to be facilitated by loci with effects restricted to individual regions such as the cerebellum (Hager et al., 2012). Nonetheless, QTL mapping is conditioned by genetic variation present in the populations used to create the crosses and the number of individuals required for detection is higher as QTL size effect decreases (Mackay et al., 2009). As such, we may miss QTL that explain variation in multiple brain components, or overall brain size, when they explain small amounts of variation, meaning we cannot fully discount additional discrete or global genetic effects influencing brain structure.

Assuming our results reflect the broad genetic architecture of variation in brain structure, why then, if brain structures have a genetic architecture that enables independent evolution, do we often observe correlated evolution among brain components across species? We suggest distributed selection pressures, rather than shared genetics, explains this pattern. For example, in fish, the diencephalon contains the preglomerular complex, a structure divided in nuclei involved in sensory, motor and limbic tasks which is similar to the thalamus in amniotes and projects neurons into the dorsal telencephalon (Mueller, 2012). The preglomerular complex receives optic tecta and retinal inputs, integrating the telencephalic ascending visual pathway (Hagio et al., 2018; Wullimann & Northcutt, 1990; Yamamoto & Ito, 2008; Zupanc, 1997). Additionally, outputs from eurydendroid cells in the cerebellum, which corresponds to the rhombencephalon, target the diencephalon (Folgueira et al., 2006; Ikenaga et al., 2002) as well as the optic tectum (Heap et al., 2013), where the visual computation required to elicit chase-avoid to objects reactions takes place (Barker & Baier, 2015; Suzuki et al., 2019). The rhombencephalon shares further functional connections with the diencephalon because the cerebellum is in turn targeted by afferent projections from the thalamic region, likely underlying visually evoked behaviours (Zompa & Dubuc, 1998). It is thus possible that the evolution of brain regions associated with sensory integration such as the preglomerular complex is associated with change elsewhere in the brain due to functional dependencies and common sources of ecological selection, as predicted under a mosaic model of brain evolution (Barton & Harvey, 2000; Montgomery et al., 2016; Avin et al., 2021).

In summary, we have provided evidence that the brain structure of the specialist lateral-line forager *A. stuartgranti* is distinct from the generalist visual-forager species *A. calliptera*, likely in part as a result of divergent sensory environments exerting selective pressures that are met with selective changes in brain components. We have also shown that this heritable variation, which is independent of allometric effects, is facilitated by a genetic architecture that allows modularity. Specifically, the majority of brain components are weakly genetically correlated despite a comparatively high degree of functional interdependence, as revealed by phenotypic correlations. In addition, some of the volumetric variation in brain components is associated with specific, independent loci with relatively small effects suggesting a distributed, polygenic basis to divergent brain morphologies. These results represent a rare test of the key predictions of major contested theories of brain evolution in a natural system.

## Data availability statement

All data and code is available from https://doi.org/10.5281/zenodo.19334348

## Acknoweldgements

We thank Dr Liz Martin-Silverstone and the XTM Facility, Paleobiology Research Group, University of Bristol for facility use and assistance, and to Dr Stephen Cross, Wolfson Bioimaging Facility for helpful discussions on image analysis. Image analysis was carried out using the computational facilities of the Advanced Computing Research Centre, University of Bristol - http://www.bristol.ac.uk/acrc/. We thank Michael Hayle for assistance with breeding and maintaining the hybrid stocks at the University of Bangor.

## CRediT Author Contributions

JM: Methodology, investigation, formal analysis, visualisation, data curation, writing the draft manuscript, reviewed and edited the final manuscript. DFRS: Formal analysis, visualisation, data curation, writing the draft manuscript, reviewed and edited the final manuscript. JE: Formal analysis, data curation, reviewed and edited the final manuscript. AH: Formal analysis, reviewed and edited the final manuscript. BF: Formal analysis, reviewed and edited the final manuscript. AM: Formal analysis, reviewed and edited the final manuscript. RD: Resources, reviewed and edited the final manuscript. GFT: Resources, reviewed and edited the final manuscript. MES: Conceptualization, funding acquisition, project administration, reviewed and edited the final manuscript. SHM: Conceptualization, funding acquisition, project administration, supervision, writing the draft manuscript, reviewed and edited the final manuscript.

## References

Abadi, M., Agarwal, A., Barham, P., Brevdo, E., Chen, Z., Citro, C., Corrado, G. S., Davis, A., Dean, J., & Devin, M. (2016). Tensorflow: Large-scale machine learning on heterogeneous distributed systems. arXiv preprint arXiv:1603.04467.

Arnold, S. J. (1992). Constraints on Phenotypic Evolution. The American Naturalist, 140, S85–S107. 10.1086/285398

Avin, S., Currie, A., & Montgomery, S. H. (2021). An agent-based model clarifies the importance of functional and developmental integration in shaping brain evolution. BMC Biology, 19(1), 97. 10.1186/s12915-021-01024-1

Barker, A. J., & Baier, H. (2015). Sensorimotor decision making in the zebrafish tectum. Current Biology, 25(21), 2804–2814. 10.1016/j.cub.2015.09.055

Barton, R. A., & Harvey, P. H. (2000). Mosaic evolution of brain structure in mammals. Nature, 405(6790), 1055–1058.

Bond, J., Roberts, E., Mochida, G. H., Hampshire, D. J., Scott, S., Askham, J. M., Springell, K., Mahadevan, M., Crow, Y. J., Markham, A. F., Walsh, C. A., & Woods, C. G. (2002). ASPM is a major determinant of cerebral cortical size. Nature Genetics, 32(2), 316–320. 10.1038/ng995

Brandstätter, R., & Kotrschal, K. (1990). Brain growth patterns in four European cyprinid fish species (Cyprinidae, Teleostei): roach (*Rutilus rutilus*), bream (*Abramis brama*), common carp (*Cyprinus carpio*) and sabre carp (*Pelecus cultratus*). Brain, Behavior and Evolution, 35(4), 195–211.

Broman, K. W., Gatti, D. M., Simecek, P., Furlotte, N. A., Prins, P., Sen, Ś., Yandell, B. S., & Churchill, G. A. (2019). R/qtl2: Software for mapping quantitative trait loci with high-dimensional data and multiparentpopulations. Genetics, 211(2), 495–502. 10.1534/genetics.118.301595

Catania, K. C. (2005). Evolution of sensory specializations in insectivores. Anat Rec A Discov Mol Cell Evol Biol, 287(1), 1038–1050. 10.1002/ar.a.20265

Chang, C. C., Chow, C. C., Tellier, L. C., Vattikuti, S., Purcell, S. M., & Lee, J. J. (2015). Second-generation PLINK: rising to the challenge of larger and richer datasets. Gigascience, 4(1), s13742-13015-10047-13748.

Danecek, P., Bonfield, J. K., Liddle, J., Marshall, J., Ohan, V., Pollard, M. O., Whitwham, A., Keane, T., McCarthy, S. A., Davies, R. M., & Li, H. (2021). Twelve years of SAMtools and BCFtools. Gigascience, 10(2). 10.1093/gigascience/giab008

Finlay, B. L., & Darlington, R. B. (1995). Linkedregularities in the development and evolution of mammalian brains. Science, 268(5217), 1578–1584. doi:10.1126/science.7777856

Finlay, B. L., Darlington, R. B., & Nicastro, N. (2001). Developmental structure in brainevolution. Behavioral and Brain Sciences, 24(2), 263–278. 10.1017/S0140525X01003958

Folgueira, M., Anadón, R., & Yáñez, J. (2006). Afferent and efferent connections of the cerebellum of a salmonid, the rainbow trout (*Oncorhynchus mykiss*): a tract-tracing study. Journal of Comparative Neurology 497(4), 542–565. 10.1002/cne.20979

Fox, J., & Weisberg, S. (2018). An R companion to applied regression. Sage publications.

Gonzalez-Voyer, A., Winberg, S., & Kolm, N. (2009a). Brain structure evolution in a basal vertebrate clade: evidence from phylogenetic comparative analysis of cichlid fishes. BMC Evolutionary Biology, 9(1), 238. 10.1186/1471-2148-9-238

Gonzalez-Voyer, A., Winberg, S., & Kolm, N. (2009b). Social fishes and single mothers: brain evolution in African cichlids. Proceedings of the Royal Society B: Biological Sciences, 276(1654), 161–167. doi:10.1098/rspb.2008.0979

Hager, R., Lu, L., Rosen, G. D., & Williams, R. W. (2012). Genetic architecture supports mosaic brain evolution and independent brain–body size regulation. Nature communications, 3(1), 1079. 10.1038/ncomms2086

Hagio, H., Sato, M., & Yamamoto, N. (2018). An ascending visual pathway to the dorsal telencephalon through the optic tectum and nucleus prethalamicus in the yellowfin goby *Acanthogobius flavimanus* (Temminck & Schlegel, 1845). Journal of Comparative Neurology, 526(10), 1733–1746. 10.1002/cne.24444

Heap, L., Goh, C.-C., Kassahn, K. S., & Scott, E. K. (2013). Cerebellar output in zebrafish: an analysis of spatial patterns and topography in eurydendroid cell projections. Frontiers in Neural Circuits, Volume 7 - 2013. 10.3389/fncir.2013.00053

Henriksen, R., Johnsson, M., Andersson, L., Jensen, P., & Wright, D. (2016). The domesticated brain: genetics of brain mass and brain structure in an avian species. Scientific Reports, 6(1), 34031. 10.1038/srep34031

Huber, R., van Staaden, M. J., Kaufman, L. S., & Liem, K. F. (1997). Microhabitat use, trophic patterns, and the evolution of brain structure in African cichlids. *Brain*, Behavior and Evolution, 50(3), 167–182.

Hutcheon, J. M., Kirsch, J. A. W., & Garland Jr, T. (2002). A comparative analysis of brain size in relation to foraging ecology and phylogeny in the chiroptera. Brain Behavior and Evolution, 60(3), 165–180. 10.1159/000065938

Ikenaga, T., Yoshida, M., & Uematsu, K. (2002). Efferent connections of the cerebellum of the Goldfish, *Carassius auratus*. Brain Behavior and Evolution, 60(1), 36–51. 10.1159/000064120

Iwaniuk, A. N., Dean, K. M., & Nelson, J. E. (2004). A mosaic pattern characterizes the evolution of the avian brain. Proceedings of the Royal Society of London. Series B: Biological Sciences, 271(suppl_4), S148-S151. doi:10.1098/rsbl.2003.0127

Joyce, D. A., Lunt, D. H., Genner, M. J., Turner, G. F., Bills, R., & Seehausen, O. (2011). Repeated colonization and hybridization in Lake Malawi cichlids. Current Biology, 21(3), R108–R109. 10.1016/j.cub.2010.11.029

Keagy, J., Braithwaite, V. A., & Boughman, J. W. (2017). Brain differences in ecologically differentiated sticklebacks. Current Zoology, 64(2), 243–250. 10.1093/cz/zox074

Konings, A. (1989). Malaŵi cichlids in their natural habitat. Cichlid Press.

Kotrschal, K., & Palzenberger, M. (1992). Neuroecology of cyprinids: comparative, quantitative histology reveals diverse brain patterns. In Environmental biology of European cyprinids (pp. 135–152). Springer.

Kruska, D. C. (2005). On the evolutionary significance of encephalization in some eutherian mammals: effects of adaptive radiation, domestication, and feralization. Brain, Behavior and Evolution, 65(2), 73–108.

Lamichhaney, S., Berglund, J., Almén, M. S., Maqbool, K., Grabherr, M., Martinez-Barrio, A., Promerová, M., Rubin, C.-J., Wang, C., Zamani, N., Grant, B. R., Grant, P. R., Webster, M. T., & Andersson, L. (2015). Evolution of Darwin’s finches and their beaks revealed by genome sequencing. Nature, 518(7539), 371–375. 10.1038/nature14181

Laughlin, S. B. (2001). Energy as a constraint on the coding and processing of sensory information. Current Opinion in Neurobiology, 11(4), 475–480. 10.1016/S0959-4388(00)00237-3

Laughlin, S. B., de Ruyter van Steveninck, R. R., & Anderson, J. C. (1998). The metabolic cost of neural information. Nature neuroscience, 1(1), 36–41.

Li, H. (2013). Aligning sequence reads, clone sequences and assembly contigs with BWA-MEM. arXiv preprint arXiv:1303.3997.

Loh, Y.-H. E., Katz, L. S., Mims, M. C., Kocher, T. D., Yi, S. V., & Streelman, J. T. (2008). Comparative analysis reveals signatures of differentiation amid genomic polymorphism in Lake Malawi cichlids. Genome Biology, 9(7), R113. 10.1186/gb-2008-9-7-r113

Loomis, C., Peuß, R., Jaggard, J. B., Wang, Y., McKinney, S. A., Raftopoulos, S. C., Raftopoulos, A., Whu, D., Green, M., McGaugh, S. E., Rohner, N., Keene, A. C., & Duboue, E. R. (2019). An adult brain atlas revealsbroad neuroanatomical changes in independently evolved populations of Mexican Cavefish. Front Neuroanat, 13, 88. 10.3389/fnana.2019.00088

Lürig, M. D., Donoughe, S., Svensson, E. I., Porto, A., & Tsuboi, M. (2021). Computer vision, machine learning, and the promise of phenomics in ecology and evolutionary biology. Frontiers in Ecology and Evolution, Volume 9 - 2021. 10.3389/fevo.2021.642774

Mackay, T. F. C., Stone, E. A., & Ayroles, J. F. (2009). The genetics of quantitative traits: challenges and prospects. Nature Reviews Genetics, 10(8), 565–577. 10.1038/nrg2612

Mahmud, M., Kaiser, M. S., McGinnity, T. M., & Hussain, A. (2021). Deep learning in mining biological data. Cognitive Computation, 13(1), 1–33. 10.1007/s12559-020-09773-x

Malinsky, M., Svardal, H., Tyers, A. M., Miska, E. A., Genner, M. J., Turner, G. F., & Durbin, R. (2018). Whole-genome sequences of Malawi cichlids reveal multiple radiations interconnected by gene flow. Nature Ecology & Evolution, 2(12), 1940–1955. 10.1038/s41559-018-0717-x

Margarido, G. R., Souza, A. P., & Garcia, A. A. (2007). OneMap: software for genetic mapping in outcrossing species. Hereditas, 144(3), 78–79. 10.1111/j.2007.0018-0661.02000.x

Meredith Jr., W. R. (1984). Quantitative Genetics. Cotton 24, 131–150. 10.2134/agronmonogr24.c5

Montgomery, S. H., Mundy, N. I., & Barton, R. A. (2016). Brain evolution and development: adaptation, allometry and constraint. Proceedings of the Royal Society B: Biological Sciences, 283(1838), 20160433. doi:10.1098/rspb.2016.0433

Moore, A. J., & Kukuk, P. F. (2002). Quantitative genetic analysis of natural populations. Nature Reviews Genetics, 3(12), 971–978. 10.1038/nrg951

Mueller, T. (2012). What is the thalamus in zebrafish? Frontiers in neuroscience, 6, 64.

Niven, J. E., & Laughlin, S. B. (2008). Energy limitation as a selective pressure on the evolution of sensory systems. Journal of Experimental Biology, 211(11), 1792–1804. 10.1242/jeb.017574

Noreikiene, K., Herczeg, G., Gonda, A., Balázs, G., Husby, A., & Merilä, J. (2015). Quantitative genetic analysis of brain size variation in sticklebacks: support for the mosaic model of brain evolution. Proceedings of the Royal Society B: Biological Sciences, 282(1810), 20151008. doi:10.1098/rspb.2015.1008

Oksanen, J., Simpson, G. L., Blanchet, F. G., Kindt, R., Legendre, P., Minchin, P. R., O’hara, R., Solymos, P., Stevens, M. H. H., & Szoecs, E. (2001). Vegan: community ecology package. 10.32614/CRAN.package.veganxf

Parsons, P. J., Bridle, J. R., Rüber, L., & Genner, M. J. (2017). Evolutionary divergence in life history traits among populations of the Lake Malawi cichlid fish *Astatotilapia calliptera*. Ecol Evol, 7(20), 8488–8506. 10.1002/ece3.3311

Pemberton, J. M. (2008). Wild pedigrees: the way forward. Proceedings of the Royal Society B: Biological Sciences, 275(1635), 613–621. 10.1098/rspb.2007.1531

Pollen, A. A., Dobberfuhl, A. P., Scace, J., Igulu, M. M., Renn, S. C., Shumway, C. A., & Hofmann, H. A. (2007). Environmental complexity and social organization sculpt the brain in Lake Tanganyikan cichlid fish. Brain Behav Evol, 70(1), 21–39. 10.1159/000101067

Rundle, H. D., & Nosil, P. (2005). Ecological speciation. Ecology letters, 8(3), 336–352.

Salzburger, W., Van Bocxlaer, B., & Cohen, A. S. (2014). Ecology and evolution of the African Great Lakes and their faunas. Annual Review of Ecology, Evolution, and Systematics, 45(Volume 45, 2014), 519-545. 10.1146/annurev-ecolsys-120213-091804

Schluter, D. (2009). Evidence for ecological speciation and its alternative. Science, 323(5915), 737–741.

Schluter, D., & McPhail, J. D. (1992). Ecological character displacement and speciation in sticklebacks. The American Naturalist, 140(1), 85–108.

Schwalbe, M. A. B., Bassett, D. K., & Webb, J. F. (2012). Feeding in the dark: lateral-line-mediated prey detection in the peacock cichlid *Aulonocara stuartgranti*. Journal of Experimental Biology, 215(12), 2060–2071. 10.1242/jeb.065920

Schwalbe, M. A. B., & Webb, J. F. (2015). The effect of light intensity on prey detection behavior in two Lake Malawi cichlids, Aulonocara stuartgranti and Tramitichromis sp. Journal of Comparative Physiology A, 201(4), 341–356. 10.1007/s00359-015-0982-y

Shumway, C. A. (2010). The evolution of complex brains and behaviors in African cichlid fishes. Current Zoology, 56(1), 144–156. 10.1093/czoolo/56.1.144

Sterling, P., & Laughlin, S. (2015). Principles of Neural Design. The MIT Press. 10.7551/mitpress/9780262028707.001.0001

Stroud, J. T., & Losos, J. B. (2016). Ecological opportunity and adaptive radiation. Annual review of ecology, evolution, and systematics, 47, 507–532. 10.1146/annurev-ecolsys-121415-032254

Suzuki, D. G., Pérez-Fernández, J., Wibble, T., Kardamakis, A. A., & Grillner, S. (2019). The role of the optic tectum for visually evoked orienting and evasive movements. Proceedings of the National Academy of Sciences, 116(30), 15272–15281. doi:10.1073/pnas.1907962116

Sylvester, J. B., Rich, C. A., Loh, Y.-H. E., van Staaden, M. J., Fraser, G. J., & Streelman, J. T. (2010). Brain diversity evolves via differences in patterning. Proceedings of the National Academy of Sciences, 107(21), 9718–9723. doi:10.1073/pnas.1000395107

Wang, J.-k., Li, Y., & Su, B. (2008). A common SNP of *MCPH1* is associated with cranial volume variation in Chinese population. Human Molecular Genetics, 17(9), 1329–1335. 10.1093/hmg/ddn021

Warton, D. I., Duursma, R. A., Falster, D. S., & Taskinen, S. (2012). smatr 3–an R package for estimation and inference about allometric lines. Methods in Ecology and Evolution, 3(2), 257–259.

Wullimann, M. F., & Northcutt, R. G. (1990). Visual and electrosensory circuits of the diencephalon in mormyrids: an evolutionary perspective. Journal of Comparative Neurology, 297(4), 537–552. 10.1002/cne.902970407

Yamamoto, N., & Ito, H. (2008). Visual, lateral line, and auditory ascending pathways to the dorsal telencephalic area through the rostrolateral region of the lateral preglomerular nucleus in cyprinids. Journal of Comparative Neurology, 508(4), 615–647. 10.1002/cne.21717

York, R.A., Byrne, A., Abdilleh, K. Patil, C., Streelman, T., Finger, T.E., & Fernald, R.D. (2019). Behavioral evolution contributes to hindbrain diversification among Lake Malawi cichlid fish. Sci Rep 9, 19994. 10.1038/s41598-019-55894-1

Zhang, J. (2023). Patterns and evolutionary consequences of pleiotropy. Annual Review of Ecology, Evolution, and Systematics, 54, 1–19. 10.1146/annurev-ecolsys-022323-083451

Zompa, I. C., & Dubuc, R. (1998). Diencephalic and mesencephalic projections to rhombencephalic reticular nuclei in lampreys. Brain Research, 802(1), 27–54. 10.1016/S0006-8993(98)00261-3

Zuiderveld, K. (1994). Contrast limited adaptive histogram equalization. In Graphics gems IV (pp. 474–485). Academic Press Professional, Inc.

Zupanc, G. K. H. (1997). The preglomerular nucleus of gymnotiform fish: relay station for conveying information between telencephalon and diencephalon. Brain Research, 761(2), 179–191. 10.1016/S0006-8993(97)00130-3

